# Omics-Based Interaction Framework – a systems model to reveal molecular drivers of synergy

**DOI:** 10.1101/2020.04.16.041350

**Authors:** Jezreel Pantaleón García, Vikram V. Kulkarni, Tanner C. Reese, Shradha Wali, Saima J. Wase, Jiexin Zhang, Ratnakar Singh, Mauricio S. Caetano, Seyed Javad Moghaddam, Faye M. Johnson, Jing Wang, Yongxing Wang, Scott E. Evans

**Affiliations:** Department of Pulmonary Medicine, University of Texas MD Anderson Cancer Center, Houston, Texas, USA 77030; MD Anderson Cancer Center UT Health Graduate School of Biomedical Sciences, Houston, Texas, USA 77030; Rice University, Houston, Texas, USA, 77005; Department of Bioinformatics and Computational Biology, University of Texas MD Anderson Cancer Center, Houston, Texas, USA 77030; Department of Thoracic, Head and Neck Medical Oncology, University of Texas MD Anderson Cancer Center, Houston, Texas, USA 77030

**Keywords:** Data integration, Inducible epithelial resistance, Multi-Omics, Pneumonia, Synergy

## Abstract

Bioactive molecule library screening strategies may empirically identify effective combination therapies. However, without a systems theory to interrogate synergistic responses, the molecular mechanisms underlying favorable drug-drug interactions remain unclear, precluding rational design of combination therapies. Here, we introduce Omics-Based Interaction Framework (OBIF) to reveal molecular drivers of synergy through integration of statistical and biological interactions in supra-additive biological responses. OBIF performs full factorial analysis of feature expression data from single vs. dual factor exposures to identify molecular clusters that reveal synergy-mediating pathways, functions and regulators. As a practical demonstration, OBIF analyzed a therapeutic dyad of immunostimulatory small molecules that induces synergistic protection against influenza A pneumonia. OBIF analysis of transcriptomic and proteomic data identified biologically relevant, unanticipated cooperation between RelA and cJun that we subsequently confirmed to be required for the synergistic antiviral protection. To demonstrate generalizability, OBIF was applied to data from a diverse array of Omics platforms and experimental conditions, successfully identifying the molecular clusters driving their synergistic responses. Hence, OBIF is a phenotype-driven systems model that supports multiplatform exploration of synergy mechanisms.

## Introduction

Superior treatment outcomes are achieved for many disease states when more than one therapeutic agent is administered (Chen *et al*, 2015; Zappasodi *et al*, 2018; Ronzitti *et al*, 2018; Han *et al*, 2019). Indeed, there are many well documented instances when the therapeutic benefit of two agents administered together substantially exceeds the benefit that would be predicted by the additive effects of the agents administered individually. Widespread availability of high throughput technologies has allowed multi-level study of complex biological responses from genome to phenome (Hasin *et al*, 2017). Yet, there remains lack of consensus regarding the appropriate analysis of statistical and biological interactions found in non-additive (i.e., antagonistic or synergistic) responses (Wei *et al*, 2018). Moreover, previously proposed strategies to analyze non-additive interactions frequently lack sufficient generalizability to study these processes outside of their home Omics platforms (Chen *et al*, 2015). Thus, while synergistic therapeutic combinations may be empirically derived from fortuitous clinical experiences or through screening of bioactive small molecule libraries, the absence of an established means to investigate these favorable drug-drug interactions ultimately precludes understanding of their underlying mechanisms. Consequently, development of a methodology to integrate the statistical and biological components of synergistic interactions in diverse Omics settings can advance the rational design of combination therapies while affording understanding of their molecular mechanisms against diseases.

Pneumonia is a major worldwide cause of death and frequently requires combination therapies (Troeger *et al*, 2017; Metlay *et al*, 2019). We have previously reported that a therapeutic dyad of immunostimulatory small molecules induces synergistic protection against a broad range of pneumonia-causing pathogens (Duggan *et al*, 2011, Cleaver *et al*, 2014, Kirkpatrick *et al*, 2018; Ware *et al*, 2019). This combination (hereafter, “Pam2-ODN”) is comprised of a Toll-like receptor (TLR) 2/6 agonist, Pam2CSK4 (“Pam2”), and a TLR 9 agonist, ODN M362 (“ODN”), that stimulate protective responses from lung epithelial cells (Cleaver *et al*, 2014). This biological response, termed inducible epithelial resistance, promotes survival benefits and microbicidal effects that significantly exceed the additive effects of the individual ligands (Duggan *et al*, 2011; Tuvim *et al*, 2012). Thus, understanding the molecular mechanisms underlying this unanticipated synergy may allow optimized manipulation of epithelial antimicrobial responses and support new generations of host-based therapeutics against infections.

In the absence of a systems theory to interrogate synergistic mechanisms (Wei *et al*, 2018), we introduce Omics-Based Interaction Framework (OBIF) to identify molecular drivers of synergy through integration of statistical and biological interactions in supra-additive biological responses. Unlike exploratory synergy models (Chen *et al*, 2015), OBIF is a phenotype-driven model (Hasin *et al*, 2017) that performs full factorial analysis (Li *et al*, 2009; Antony, 2014; Das *et al*, 2018) of feature expression data from single vs. dual factor exposures to identify molecular clusters that reveal synergy-mediating pathways, functions and regulators. To demonstrate the utility of OBIF, we applied this strategy to multi-Omics experimental data from epithelial cells exposed to Pam2-ODN to identify biologically relevant, unanticipated cooperative signaling events that we subsequently confirmed to be required for the synergistic pneumonia protection. Then, to demonstrate generalizability, OBIF was applied to datasets from diverse types of Omics platforms and experimental models, successfully identifying molecular clusters driving their synergistic responses.

## Results

### Synergistic Pam2-ODN-induced epithelial resistance against pneumonia

Our laboratory’s interest in synergistic interactions arises from our experience investigating single vs. dual immunostimulatory treatments to prevent pneumonia (Duggan *et al*, 2011; Tuvim *et al*, 2012; Cleaver *et al*, 2014; Kirkpatrick *et al*, 2018; Ware *et al*, 2019). As a demonstrative example, data are presented here from influenza A virus (IAV) challenges of different models following pretreatment with Pam2 alone, ODN alone or the Pam2-ODN combination. When mice are challenged with IAV 24 h after the indicated inhaled treatments, we observed little increase in survival after the individual treatments compared to sham-treated control mice, whereas mice treated with the Pam2-ODN combination demonstrated profound antiviral protection (Figure 1A). Similarly, when isolated mouse lung epithelial (MLE-15) cells were challenged with IAV 4 h after pretreatment with the individual ligands, we observed no significant reductions in the viral burden relative to PBS treated cells. However, cells pretreated with Pam2-ODN showed a substantial reduction in viral nucleoprotein (NP) gene expression as assessed by qPCR relative to host 18s gene (Figure 1B). Comparing the effect of dual ligand treatment (E_AB_) to the expected response additivity of the individual ligand treatments (E_A_ + E_B_) (Foucquier *et al*, 2015) reveals supra-additive effects on both *in vivo* survival benefits and *in vitro* viral clearance (Figure 1C). To better understand the molecular mechanisms driving such unanticipated synergy, we developed OBIF as a phenotype-driven model (Hasin *et al*, 2017) to understand favorable drug-drug interactions mediating synergistic responses and outcomes.

**Figure 1.**
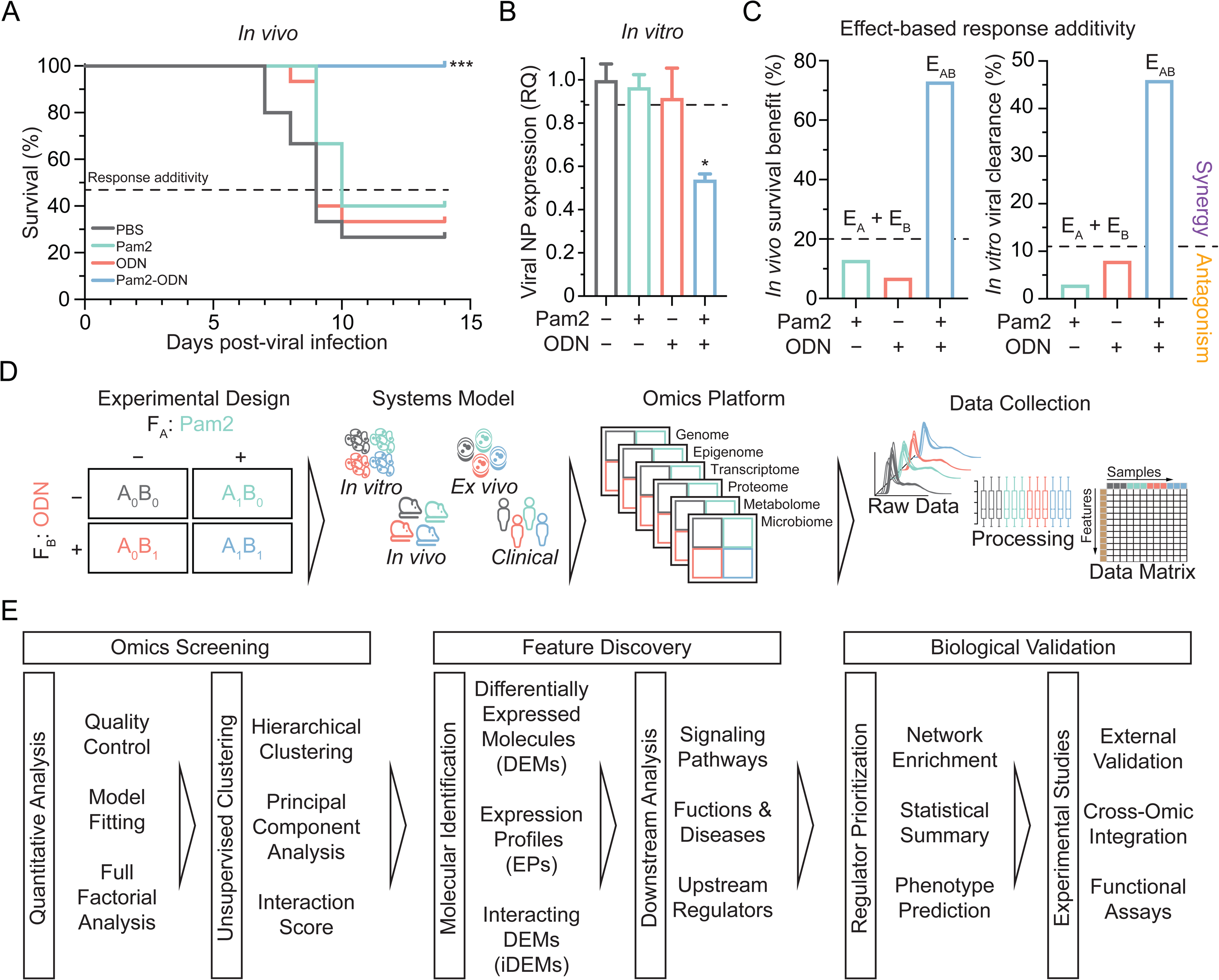
Omics-Based Interaction Framework: phenotype-driven synergy modeling and framework overview. (A) Mouse survival of influenza A challenge following the indicated pretreatments. Dashed line indicates additive effect of single ligand treatments over PBS. n = 15 mice/condition. (B) Virus burden of isolated mouse lung epithelial cells after influenza A challenge following the indicated pretreatments. RQ, relative quantification of viral nucleoprotein (NP) expression to host 18s. n = 6 samples/condition. (C) Plot of response additivity from antiviral responses in panels A (left) and B (right). Synergistic effects are reflected by E_AB_ greater than the expected linear sum (E_A_ + E_B_, dashed line) of individual ligand effects, antagonistic effects are observed when E_AB_ < E_A_ + E_B_. (D) Generic Omics workflow for phenotype-driven synergy modeling using a 22 experimental design. (E) Overview of OBIF, including (i) Omics screening of features in a data matrix, (ii) discovery of feature clusters of molecular drivers, and (iii) experimental validation of biologically relevant synergy regulators. * P < 0.05 compared to either condition, *** P < 0.0005 compared to either condition.

### Development of a systems synergy model from experimental Omics data

To formally test whether the effect of dual factors (F_AB_: Pam2-ODN) is greater than the expected linear sum of its individual factors (F_A_: Pam2; F_B_: ODN), an initial 2-level 2-factor (2^2^) factorial design is required (Slinker, 1998; Foucquier *et al*, 2015) (Figure 1D). Our strategy adapts the traditional analysis of variance (ANOVA) approach into a model that links the empirical analysis of synergy (Slinker, 1998; Foucquier *et al*, 2015) with the high-throughput capacity and high-dimensionality of Omics datasets (Coral *et al*, 2017; Bardini *et al*, 2017). As summarized in Figure 1E, OBIF integrates statistical and biological interactions in Omics data matrices from single vs. dual factor exposures, leveraging Omics screening to promote discovery of the molecular drivers of synergy, and facilitating the biological validation of synergy regulators. The analytical pipeline is freely available as an R package at GitHub (www.github.com/evanslaboratory/OBIF). Naturally, the experimental validation components must be tailored to the individual tools and characteristics of the biological responses being studied.

### Differentially expressed molecules reveal synergy-specific pathways

To investigate the mechanisms underlying Pam2-ODN synergy, we used OBIF to re-analyze previously published (Data Ref: Tuvim *et al*, 2014) lung homogenate transcriptomic data from mice inhalationally treated with single vs. dual ligands (GSE28994). After model fitting for feature expression (Figure EV1), this analysis identifies 3456 features as differentially expressed molecules (DEMs) 2 h after treatment with Pam2, 2941 DEMs after ODN treatment, and 3138 DEMs after treatment with Pam2-ODN (Figure 2A). Despite the fact that 52% (1617/3138) of DEMs were shared by Pam2-ODN and the individual ligands, enrichment analysis using IPA software revealed an overrepresentation of 12 canonical cellular immune response and cytokine signaling pathways that were activated by Pam2-ODN but not by either or both single ligands (Figure 2B). Of these, NF-κB signaling was the most enriched signaling pathway by Pam2-ODN treatment. Although unsupervised hierarchical clustering consistently segregated the treatment groups (Figure 2C), this approach alone did not reveal distinctive gene clusters to explain the synergistic response, likely due to the 52% redundancy of DEMs between groups.

**Figure 2.**
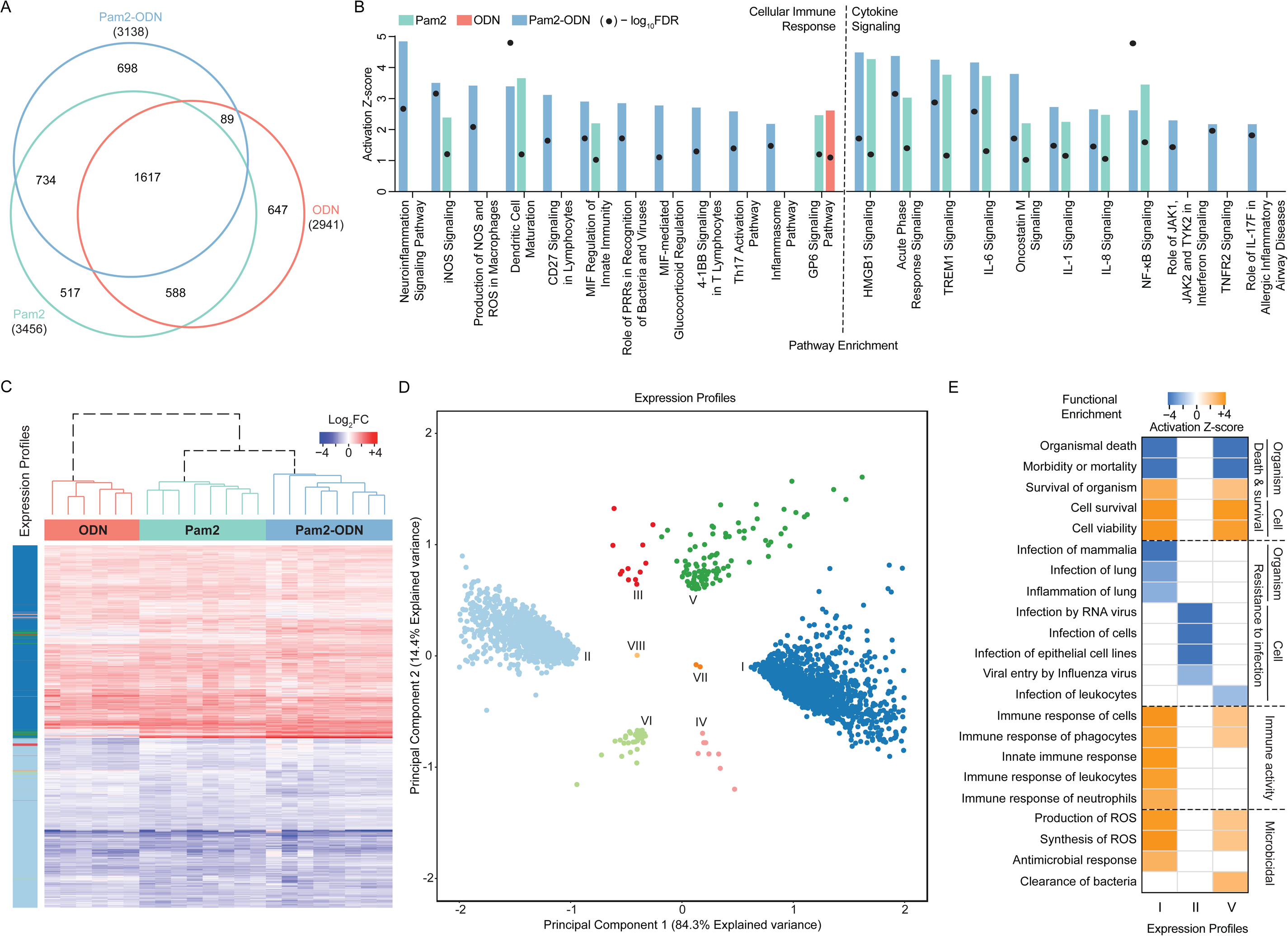
Differentially expressed molecules and expression profiles reveal synergy-mediating pathways and functions. (A) Euler diagram of differentially expressed molecules following single or dual treatment in mouse lung homogenates. (B) Most overrepresented activated canonical pathways after IPA enrichment of DEMs. (C) Heatmap of expression values of DEMs in A with expression profiles shown per feature (rows). (D) Principal component analysis of Pam2-ODN DEMs identified by expression profiles. (E) Top activated (orange) and inhibited (blue) diseases and functions after IPA enrichment of expression profiles. FC, fold change.

### Expression profiles summarize biological interactions and disentangle effectors of synergistic functions

Rather than relying on potentially redundant DEM clusters, OBIF classifies dual factor-induced DEMs into eight expression profiles (EPs) that characterize cooperative and competitive biological interactions of individual factors (Table 1). EPs are defined by expression directionality (up- or down-regulation) of individual features and are not biased by the DEM expression analysis. Cooperative EPs have accordant expression directionality induced by F_A_ and F_B_, while competitive EPs have opposite directionalities induced by F_A_ and F_B_. Among the cooperative EPs, concordant profiles result when F_AB_ directionality corresponds with the single factor effects (EPs I and II), and discordant profiles occur when F_AB_ directionality opposes the single factors (III and IV). Alternatively, among the competitive EPs, factor-dominant profiles are defined by F_AB_ directionality correspondence with one factor (F_A_ dominant, V and VI; F_B_ dominant, VII and VIII). Principal component analysis (Figure 2D) demonstrates that concordant EPs (I and II) were the most abundant in our dataset, followed by Pam2-dominant profiles (V and VI). This abundance of EPs I and II better emphasizes the cooperative effects of both factors than does conventional DEM clustering alone. In particular, the contribution of ODN to the synergistic combination might otherwise be overlooked by DEM analysis, as it induces enrichment of far fewer signaling pathways (Figure 2B) and has a greater clustering distance from Pam2-ODN samples (Figure 2C).

**Table 1.**
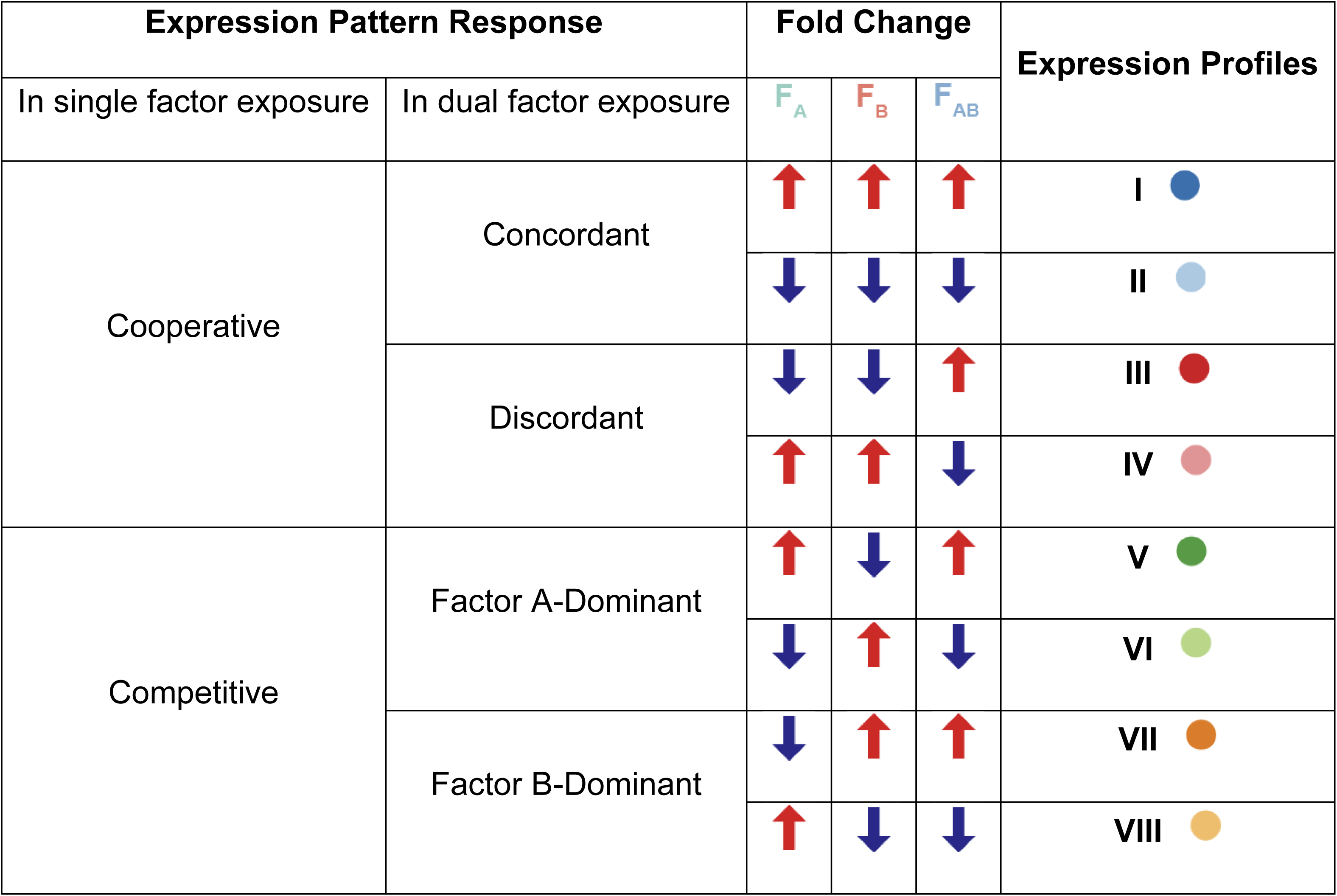
Expression profiles depict biological interactions during dual factor exposure.

| Expression Pattern Response |  | Fold Change |  |  | Expression Profiles |
| --- | --- | --- | --- | --- | --- |
| In single factor exposure | In dual factor exposure | F<sub>A</sub> | F<sub>B</sub> | F<sub>AB</sub> |  |
| Cooperative | Concordant | ↑ | ↑ | ↑ | I ● |
|  |  | ↓ | ↓ | ↓ | II ● |
|  | Discordant | ↓ | ↓ | ↑ | III ● |
|  |  | ↑ | ↑ | ↓ | IV ● |
| Competitive | Factor A-Dominant | ↑ | ↓ | ↑ | V ● |
|  |  | ↓ | ↑ | ↓ | VI ● |
|  | Factor B-Dominant | ↓ | ↑ | ↑ | VII ● |
|  |  | ↑ | ↓ | ↓ | VIII ● |

Notably, enrichment analysis reveals that molecular effectors clustered by EPs correspond with Pam2-ODN-induced functions (Figure 2E), suggesting a biological basis for the synergy. Specifically, we found that features in profiles I and V contributed to host survival functions, immune activity and microbicidal activity. Considered from an organizational perspective, induction of resistance to infection at the organismal level correlated with features in profile I, at the cellular level with profile II, and by leukocytes with profile V.

### Factorial effects analysis integrates biological and statistical interactions in EPs

Analysis of factorial effects in a data matrix from single vs. dual factor exposures can statistically differentiate whether stochastic feature expression in a combination is correlated with the effect of an individual factor (simple main effect, SME) or their influence on each other (interaction effect) (Li *et al*, 2013; Mihret *et al*, 2014; Zhang *et al*, 2017). Based on this principle, OBIF performs full factorial analysis through paired comparisons of calculated β coefficients in each condition to determine statistical relationships (Hassall *et al*, 2018) and discover significant main effects during expression analysis and multi-factor effects (SMEs and interaction effect) from contrast (Mee, 2009) analysis (Figure 3A). Using this approach, main effects determined significant DEMs per condition, while multi-factor effects explained whether Pam2-ODN DEMs and EPs resulted from SMEs and/or an interaction of individual ligands (Figure 3B). This analysis showed that most features in concordant profiles (I and II) are influenced by at least one multi-factor effect, while all features in discordant profiles (III and IV) are influenced by all multi-factor effects simultaneously. Not surprisingly, Pam2-dominant (V and VI) and ODN-dominant (VII and VIII) expression mainly results from their respective SMEs. This analysis also revealed that 67% (2116/3138) of Pam2-ODN DEMs are driven by the interaction effect of Pam2 and ODN as interacting DEMs (iDEMs) (Figure 3C). Thus, OBIF reconciled the biological interactions from EPs with the statistical interactions from multi-factor effects of Pam2-ODN.

**Figure 3.**
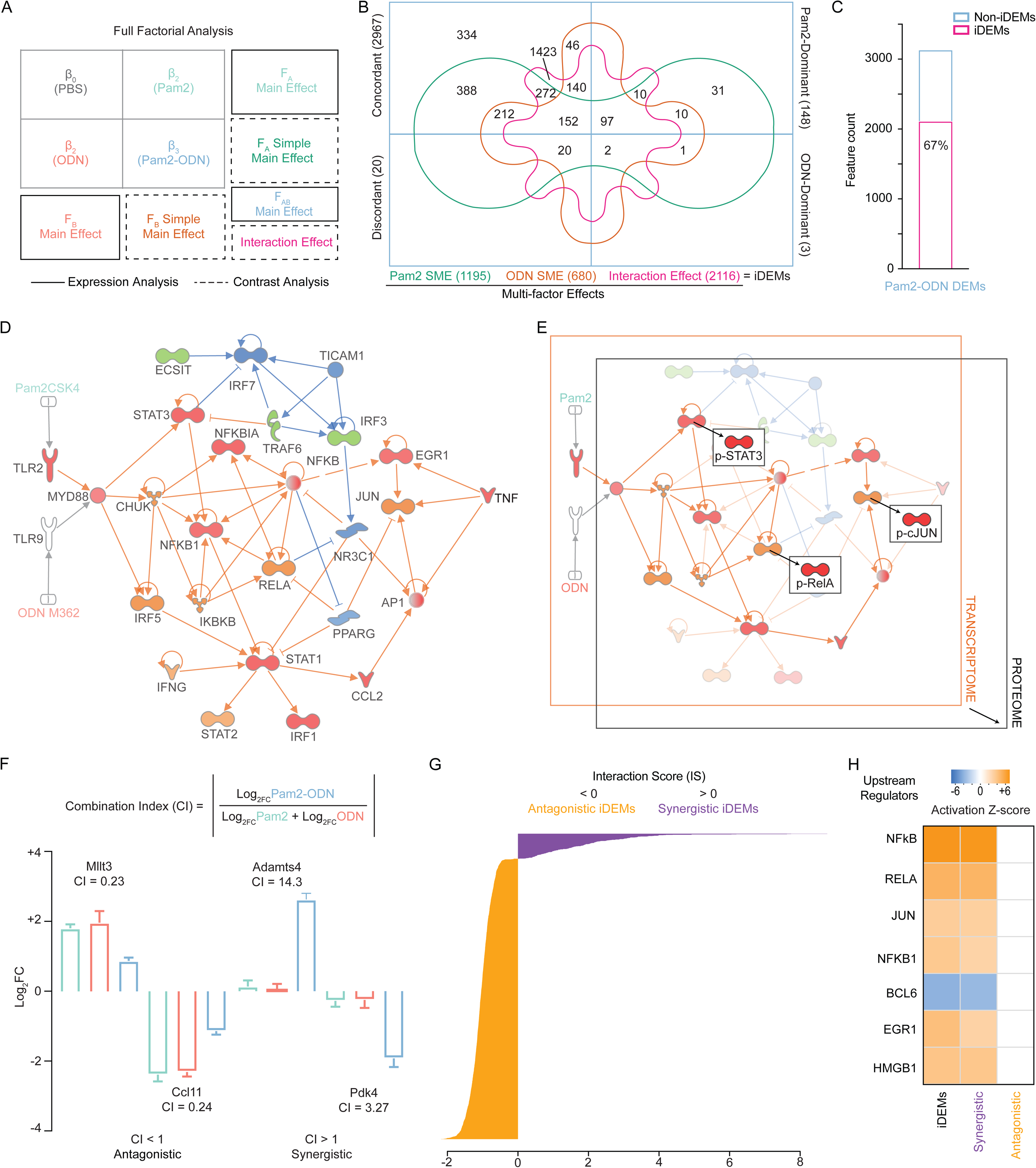
Full factorial analysis reveals regulatory networks and molecular drivers of synergy. (A) Scheme of full factorial analysis performed by OBIF from β coefficients. (B) Venn diagram of Pam2-ODN DEMs correlated by expression profiles classes and multi-factor effects. (C) Feature count and percentage of Pam2-ODN iDEMs and non-iDEMs in B. (D) Network analysis of activated (orange) or inhibited (blue), and up-regulated (red) or down-regulated (green) upstream regulators of Pam2-ODN after IPA enrichment of SME. (E) Cross-Omics validation of regulators in D. Differentially expressed phospho-signaling molecules were identified with OBIF from a reverse-phase protein array in human lung epithelial cells. (F) Non-additive feature expression assessed by combination index (CI). Representative genes and their CI values are shown. (G) Interaction score (IS) of iDEMs, reflecting antagonistic (IS < 0) and synergistic (IS > 0) features. (H) Top activated (orange) or inhibited (blue) transcriptional regulators after IPA enrichment of iDEMs. DEMs, differentially expressed molecules. iDEMs, interacting DEMs. SME, simple-main effects.

### SMEs accurately reproduce the regulatory network of combined exposures

Downstream analyses of SMEs have the capacity to discern the contributing roles of individual factors to a combination treatment (Hassall *et al*, 2018). Hence, Pam2-ODN DEMs with significant SMEs were used for network analysis of upstream regulators that are activated (orange) or inhibited (blue) and up-regulated (red) or down-regulated (green) (Figure 3D). Similar to our previous findings with DEMs (Figure 2B), transcription factors from many pathways were involved, though NF-κB family members remained central elements of this network. Demonstrating the cross-Omics function of OBIF, a parallel analysis of reverse-phase protein array (RPPA) data from single- or dual-treated human lung epithelial cells identified the top phospho-signaling DEMs (Figure EV2), and cross-validated STAT3, RelA and cJun as transcriptional units involved in the Pam2-ODN signaling network (Figure 3E).

### iDEMs identify non-additive features and synergy regulators

Non-additivity results from strong interaction effects between two factors in a combination and gives rise to synergistic or antagonistic responses (Slinker et al, 1998; Geary et al, 2013). iDEMs integrate this principle during feature selection based on significant interaction effects between factors, allowing quantification of synergistic and antagonistic expression in a narrower set of differentially expressed features. OBIF builds on previous definitions of the combination index (CI) (Foucquier *et al*, 2015; Goldstein *et al*, 2017) to fit the values of feature expression:

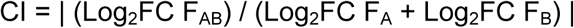

where CI is the absolute ratio of the log2 fold change of Pam-ODN-induced DEMs (F_AB_) and the additivity threshold of Pam2 (F_A_) and ODN (F_B_), allowing identification of both antagonistic (CI < 1) or synergistic (CI > 1) features (Figure 3F). A log2 transformation of the CI then yields an interaction score (IS) that quantifies the effect size of non-additive expression relative to the additivity threshold, and can be applied to both antagonistic (IS < 0) and synergistic (IS > 0) iDEMs (Figure 3G). This allows more focused enrichment analysis, in this case supporting NF-κB/RelA and AP-1/cJun as key transcriptional upstream regulators of Pam2-ODN’s interaction effect and synergistic expression (Figure 3H).

### Experimental validation of molecular regulators of Pam2-ODN synergy

Prompted by the foregoing results, we tested whether RelA and cJun were biologically relevant synergy regulators of Pam2-ODN-induced epithelial resistance. The DNA-binding activity of NF-κB and AP-1 subunits in isolated human bronchial epithelial cells (HBEC-3kt) after stimulation with Pam2-ODN confirmed that RelA and cJun activation was strongly increased after 15 minutes of treatment without significant contribution of other family members (Figure 4A, Figure EV3A). Indeed, RelA and cJun exhibited surprisingly similar activation kinetics after Pam2-ODN treatment, further supporting cooperation or coordination (Figure 4B). Investigating this co-activation of non-redundant transcriptional families, single-cell nuclear translocation of canonical p50/RelA and cFos/cJun dimers in HBEC-3kt was assessed by imaging flow cytometry. We found that all transcriptional subunits exhibited an increased nuclear translocation (similarity score > 2) after 15 minutes of Pam2-ODN treatment relative to the PBS-treated cells (Figure 4C). However, neither Pam2 nor ODN alone induced the same magnitude of nuclear translocation, whether assessed by similarity scores (Rd value) or by the percentage of translocated cells (Figure 4D) relative to PBS treated cells.

**Figure 4.**
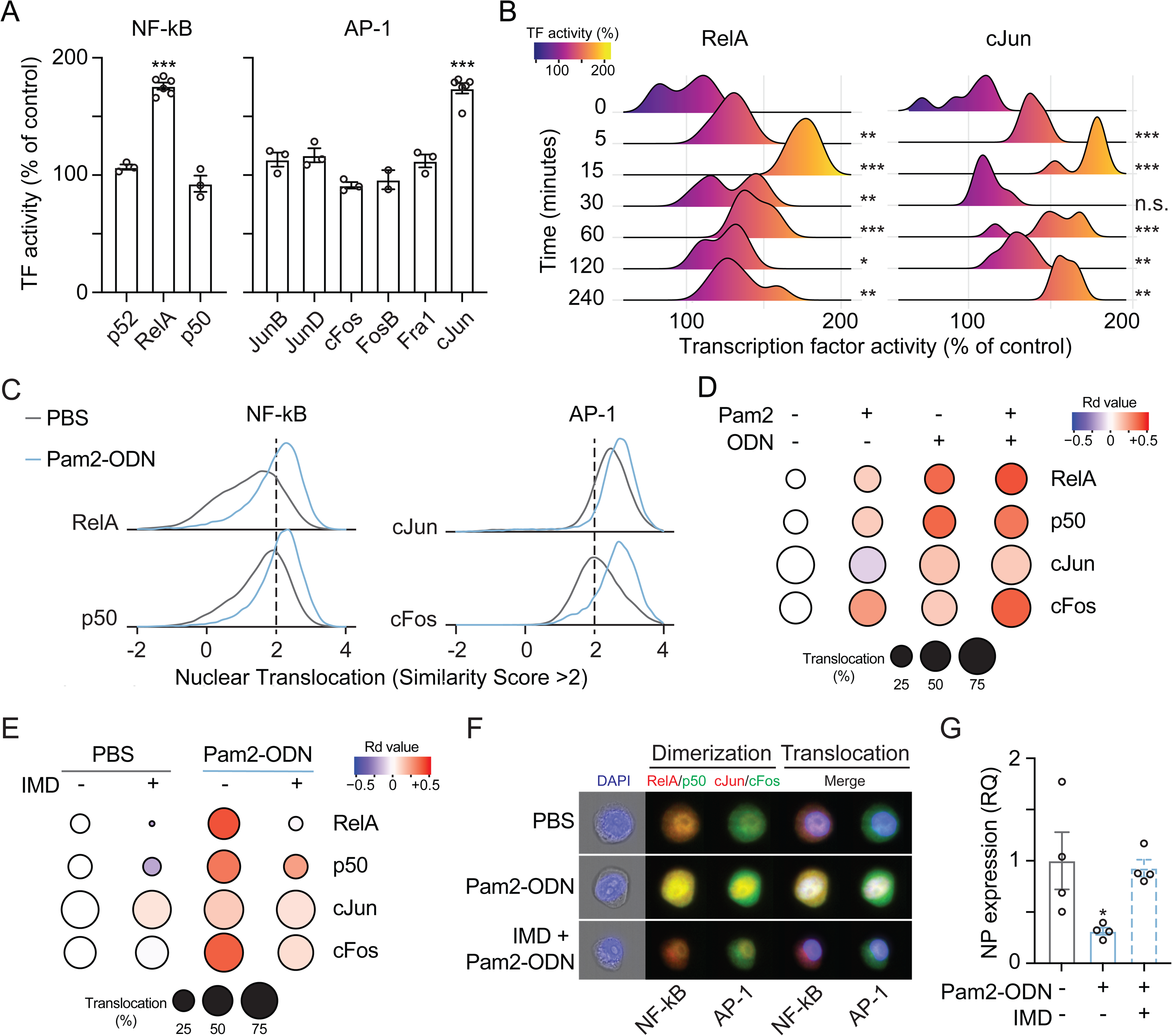
Regulators identified with OBIF uncovered cooperation between RelA and cJun that is required for synergistic antiviral protection. (A) Transcription factor activity of NF-κB and AP-1 subunits 15 min after treatment of human lung epithelial cells with Pam2-ODN. n = 3-6 samples/condition. (B) RelA and cJun activity at indicated times after Pam2-ODN treatment. n = 6 samples/condition. (C) Nuclear translocation scores of NF-κB and AP-1 heterodimers after Pam2-ODN treatment. Shown as normalized frequency of similarity score per condition. (D) NF-κB and AP-1 subunit nuclear translocation in C (increased, red; decreased, blue) per condition. (E) NF-κB and AP-1 subunit nuclear translocation with or without NF-κB inhibition by IMD-0345. (F) Representative imaging flow cytometry images of hetero-dimerization and nuclear translocation of NF-κB and AP-1 in D and E. (G) Virus burden of mouse lung epithelial cells challenged with influenza A with or without NF-κB inhibition. n = 4 samples/condition. *, P < 0.05; **, P < 0.005; ***, P < 0.0005 compared to baseline.

### Discovery of novel NF-κB and AP-1 cooperation required for antiviral protection

To differentiate transcriptional cooperation from coincidental transcription factor activation after Pam2-ODN treatment, we assessed the Pam2-ODN-induced nuclear co-translocation of NF-κB and AP-1 complexes in the presence or absence of NF-κB inhibitor IMD-0354 (IMD). As expected, pre-treatment with IMD alone reduced the Rd Value and percentage of translocated cells for RelA and p50 without significantly modifying the percentage of translocation for cJun and cFos. However, NF-κB inhibition with IMD also unexpectedly reduced the Pam2-ODN-induced similarity score shifts and nuclear translocation of AP-1 subunits, particularly of cFos (Figure 4E). This indicates that NF-κB inhibition impaired Pam2-ODN-induced AP-1 nuclear translocation, confirming the cooperative regulation of these two non-overlapping signaling pathways. Representative images shown in Figure 4F demonstrate that inhibition with IMD reduced Pam2-ODN-induced heterodimerization and nuclear translocation of NF-κB and AP-1 complexes. Further, we confirmed that disruption of this transcriptional cooperation was sufficient to impair the inducible viral burden reduction seen with Pam2-ODN (Figure 4G).

### Application of OBIF across multiple platforms and conditions

To demonstrate its generalizability, we used OBIF to analyze synergistic regulators in datasets derived from microarray, RNA-seq, RPPA and mass spectrometry-based metabolomics investigations of diverse factor classes and biological systems that demonstrate synergistic biological outcomes (Data Ref: Tuvim *et al*, 2014; Data Ref: Caetano *et al*, 2018; Data Ref: Singh *et al*, 2019; Data Ref: Han *et al*, 2019). As a preliminary step before full factorial analysis of individual features, OBIF performs an interaction analysis between the two factors of interest using a two-way ANOVA model to represent the impact of factorial effects at the whole “-ome” level. This statistical summary shows the effects of individual factors and interactions through interaction plots and statistical significance calculations (Figure 5A). This provides adjusted R^2^ and F-statistic p-values of the two-way ANOVA that allow evaluation of improved model fitness (Figure EV4 A) and detection of interaction terms (Figure EV4 B) within a dataset. After confirming adequate model fitness (i.e. adjusted R^2^ > 0.5, F-test < 0.05), full factorial analysis on scaled data from targeted or non-targeted platforms identifies DEMs (Figure 5B) from individual features with an increased discriminatory power for interaction effects (Figure EV4C). EPs then represent the biological interactions of dual factor DEMs regardless of their factor classes (Figure 5C). Contrast analysis is then applied to more adequately retrieve and classify iDEMs (Figure EV4D) and interaction scores are calculated in a uniform scale whether the original data contained continuous or count-based expression values (Figure 5D). Finally, OBIF visually summarizes the results of full factorial analysis in a Circos plot to easily identify molecular drivers of synergy from the co-expressed features, DEMs, log_2_FC, EPs, multi-factor effects and iDEMs with their interaction score (Figure 5E).

**Figure 5.**
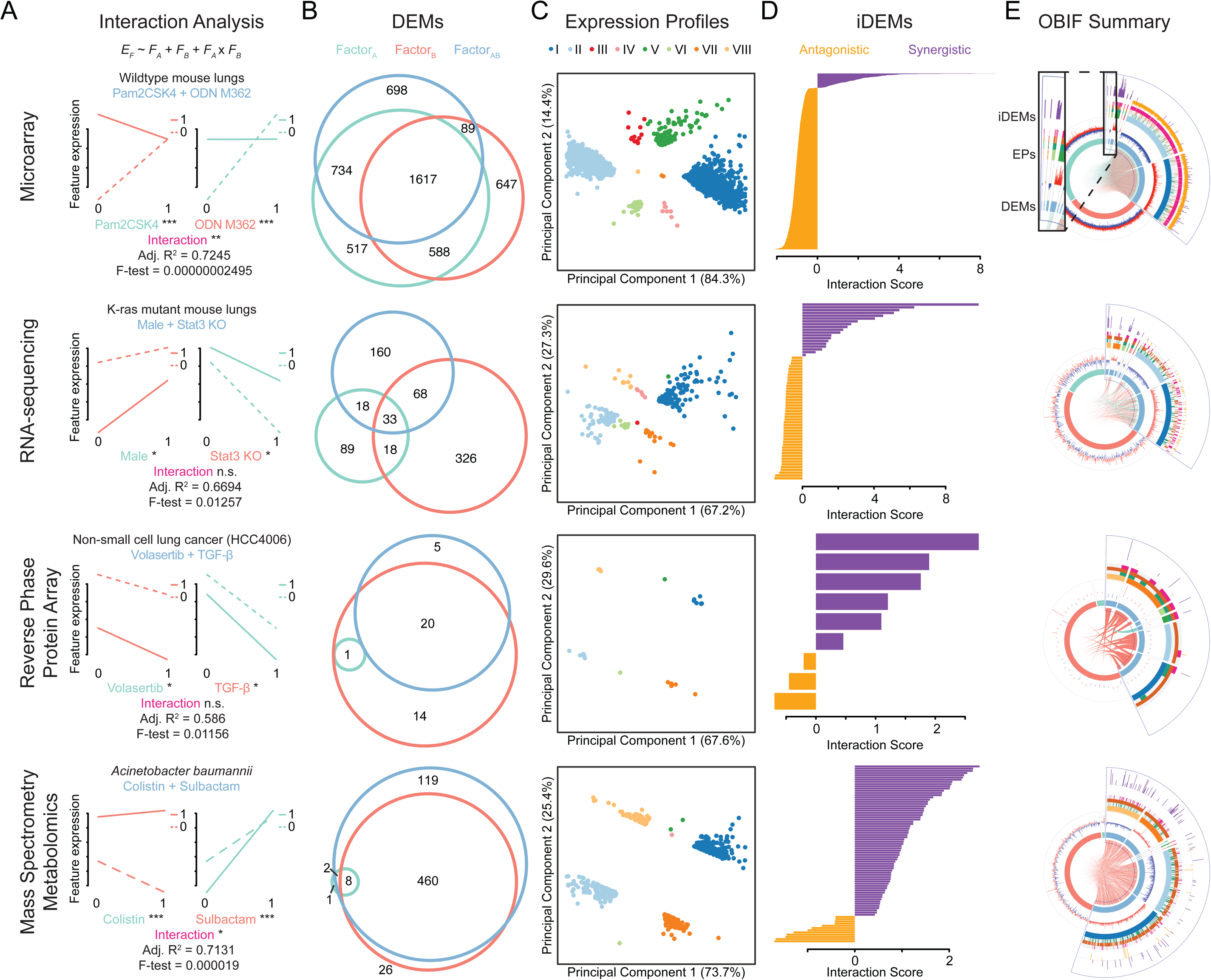
OBIF reveals molecular drivers of synergy across platforms, factor classes and experimental systems. (A) Interaction analysis of factorial effects at the whole “-ome” level, demonstrating interaction plots, coefficient significance and quality of model fitness per platform. (B) Euler diagram of DEMs identified in A. (C) Principal component analysis of dual factor DEMs in B clustered by EPs. (D) Interaction scores of iDEMs in C. (E) Visual summary of molecular drivers of synergy in B-D plotted including DEMs, EPs and iDEMs. E_F_, feature expression; F_A_, factor A; F_B_, factor B; F_AB_, factor AB; FC, fold change; DEMs, differentially expressed molecules; EPs, expression profiles; iDEMs, interacting DEMs.

## Discussion

Synergistic and antagonistic interactions are common in nature and frequently promote efficacy of therapeutic interventions (Chen *et al*, 2015; Ronzitti *et al*, 2018; Wei *et al*, 2018; Zappasodi *et al*, 2018; Han *et al*, 2019). While synergy quantification methods from dose-response data, combinatorial screening of molecule libraries, and other predictive exploration models may suggest potentially synergistic conditions or treatments, they do not provide substantive insights into the molecular mechanisms underlying synergy (Chen *et al*, 2015). Thus, synergy-mediating pathways cannot be strategically targeted in rational drug development.

Our interest in synergy arose from our observations of the strikingly synergistic interactions of one such empirically derived combination, Pam2-ODN. While we could easily quantify the superiority of protection conferred by the dual treatment, in the absence of a systems theory to interrogate synergistic mechanisms (Chen *et al*, 2015; Wei *et al*, 2018), we were limited in our capacity to use available Omics datasets to deduce the mechanisms mediating the synergy. This is important because, although this lack of mechanistic understanding does not limit the utility of the current combination, it precludes development of next generation interventions that more precisely (perhaps, more efficaciously) target the synergy-driving pathways with fewer off-target (potentially toxic) effects. In contrast to models that predict possible synergy, OBIF was developed with the explicit intent to investigate established synergistic events. As such, it is inherently a phenotype-driven model that performs full factorial analysis on feature expression data from single vs. dual factor exposures to identify molecular clusters that reveal synergy-mediating pathways, functions and regulators.

Using Pam2-ODN datasets as demonstrative examples, OBIF identified unanticipated transcriptional cooperation between non-redundant transcription factors, RelA and cJun, as a molecular mechanism of inducible synergistic protection against IAV. Thus, by facilitating understanding of combined factor exposures in terms of the individual components, a computational discovery facilitated experimental validation of a discrete, novel mediator of a non-additive biological response. Perhaps as importantly, the computational analyses were accomplished by integration of data from different Omics platforms, different specimen types, and even different host species.

Unlike most 2^2^ designs, OBIF dissects factorial effects of dual factor exposures through full factorial analysis of feature expression data in a single unsupervised step. This allows simultaneous identification of DEMs directly from main effects of single or dual factors, overcoming pairwise comparisons to control and repetitive analysis of each condition. While this simultaneous identification of DEMs can be performed also with a mixed-effect model, we showed how this approach is suboptimal to detect interaction effects at the level of individual features and iDEM selection when compared with full factorial analysis. Additionally, clustering by DEMs, EPs and iDEMs improves the specificity of enrichment analysis to disentangle the signaling pathways, functions and regulators of this synergistic combination and to capture their specific driving features. Further, quantification of multi-factor effects (SMEs and interaction effects) reveals whether particular features, molecular clusters or functions enriched by synergistic combinations are the result of individual factors or their crosstalk.

These statistical relationships have biological analogues that are integrated by OBIF in the EP definitions. In fact, profiles I and II rescued the underrepresentation of ODN observed in distance-based clustering and enrichment analysis. Further, iDEMs derived from features with significant interaction effects allow focusing discovery on synergy regulators and the calculation of interaction scores allows quantification of their non-additive expression. Thus, unlike most systems models of synergy, OBIF facilitates integrative analyses of biological and statistical interactions that are easily discoverable and interpretable through molecular clusters representing the complex dynamics of synergistic combinations.

OBIF is available as an open-source R package with a semi-automated pipeline to facilitate its broad application to unscaled original data from various Omics platforms, factor classes and biological systems. We have shown that OBIF can be fitted to perform full factorial analysis and that it adequately identifies DEMs, EPs, iDEMs and their attendant values and scores to promote discovery of molecular drivers of synergy in multiple, diverse datasets.

In summary, OBIF provides a phenotype-driven systems biology model that allows multiplatform dissection of molecular drivers of synergy. And, we encourage the application of OBIF to provide holistic understanding in research fields where greater-than-additive beneficial combinations remain understudied.

## Materials and Methods

### Reagents and Tools Table

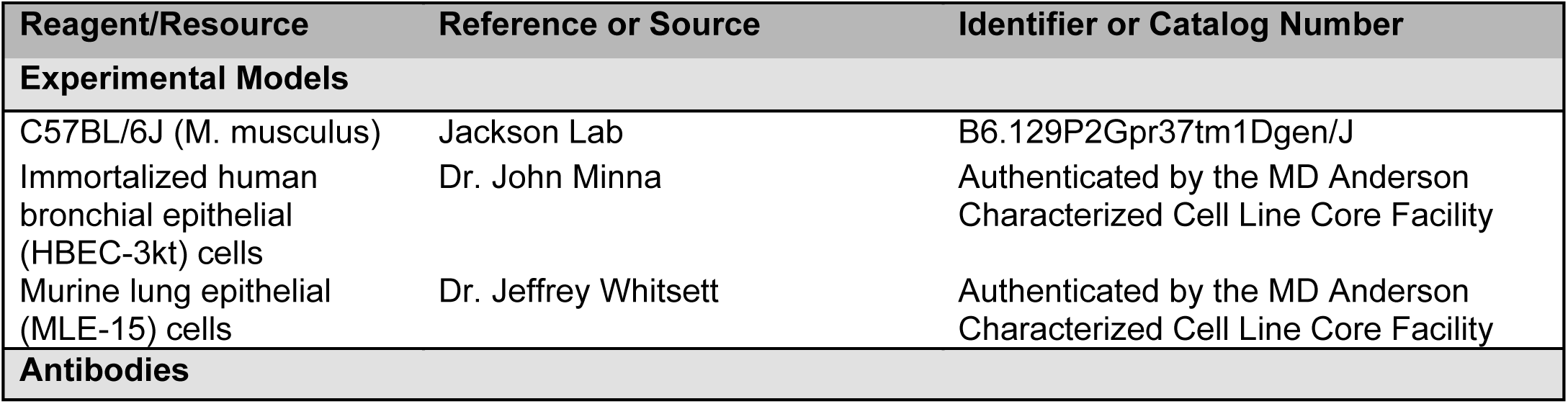
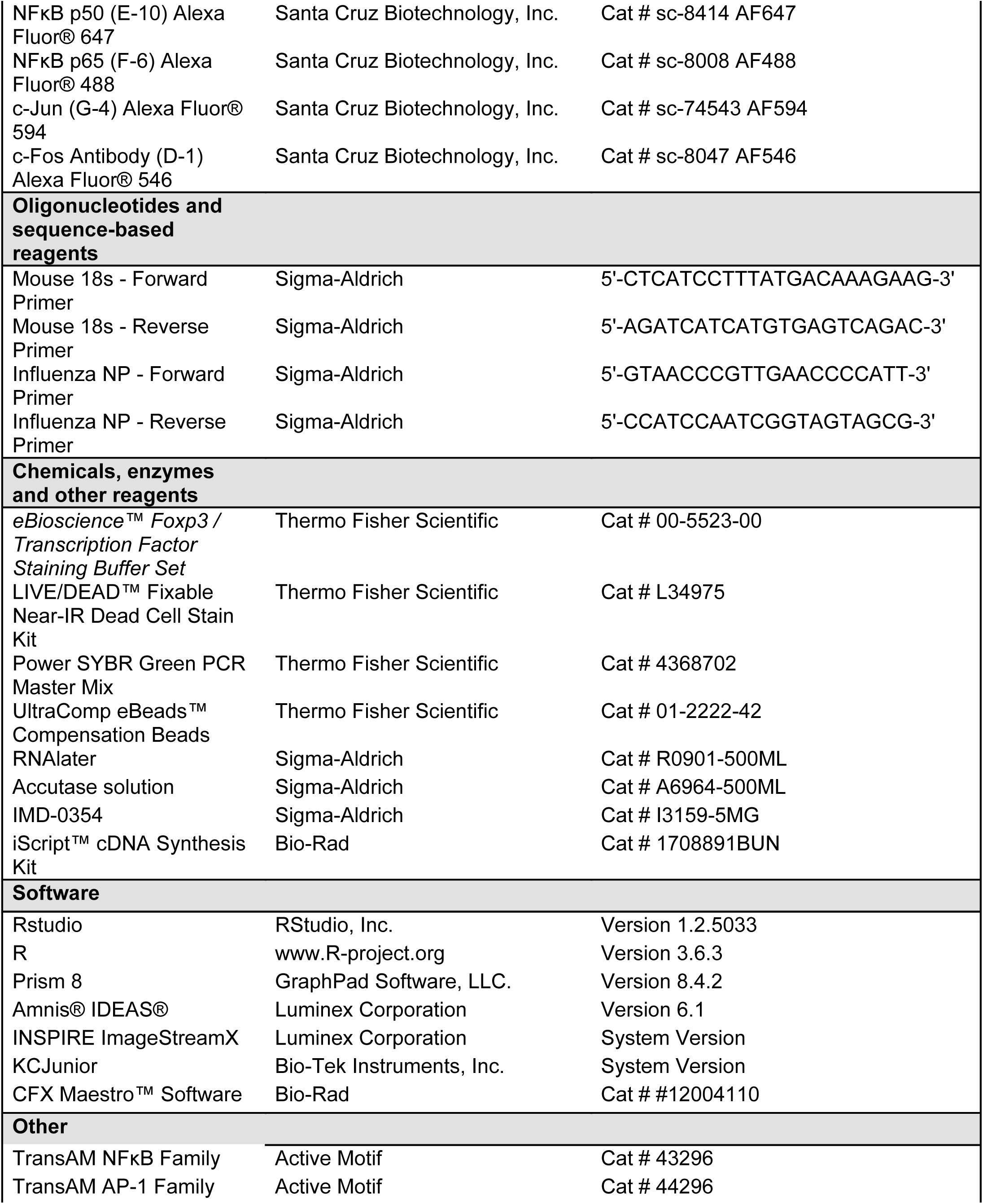
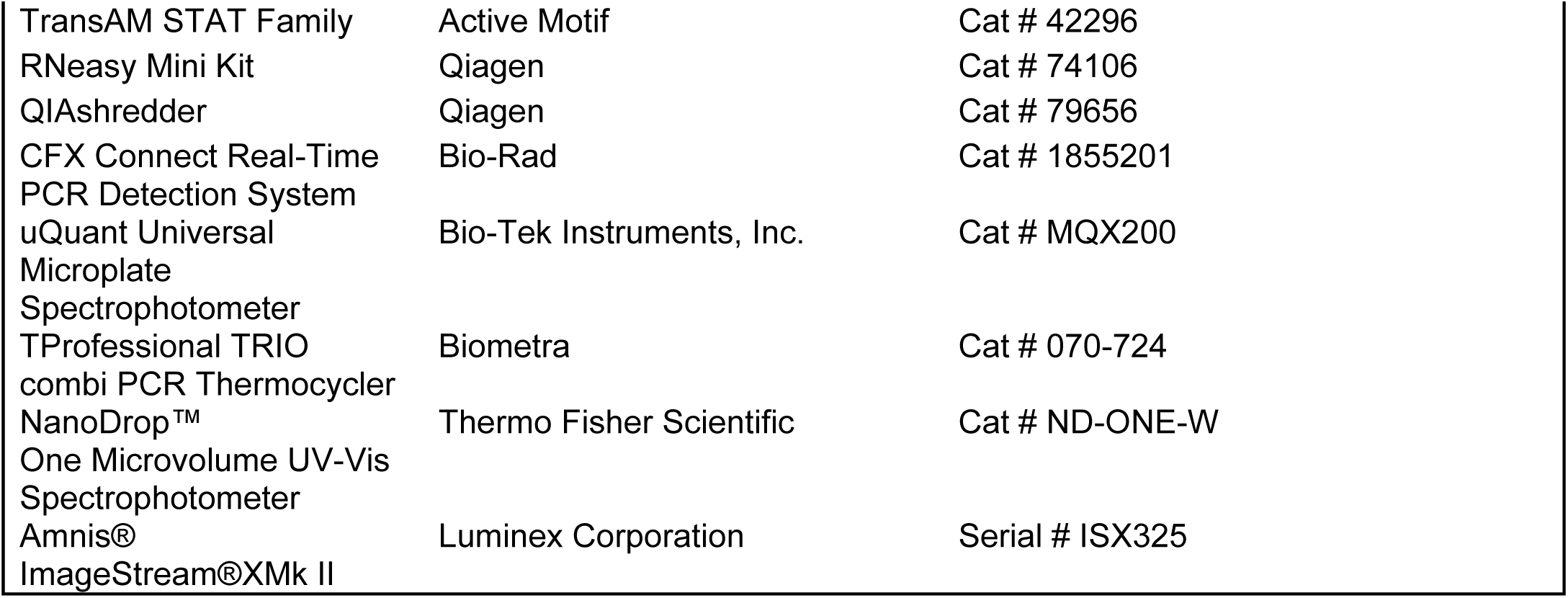

## Methods and Protocols

### Experimental Models

#### Animals

All mouse experiments were performed with 6-10 week old C57BL/6J mice of a single sex in accordance with the Institutional Animal Care and Use of Committee of The University of Texas MD Anderson Cancer Center, protocol 00000907-RN01.

#### Cell culture

Immortalized human bronchial epithelial (HBEC-3kt) cells were kindly provided by Dr. John Minna. HBEC-3kt cells were cultured in keratinocyte serum-free media (KSFM) supplemented with human epidermal growth factor and bovine pituitary extract. Murine lung epithelial (MLE-15) cells were kindly provided by Dr. Jeffrey Whitsett. The cell lines used were authenticated by the MD Anderson Characterized Cell Line Core Facility. MLE-15 cells were cultured in RPMI supplemented with 10% fetal bovine serum. Cultures were maintained in the presence of penicillin and streptomycin.

### Exposure to TLR ligands

S-[2,3-bis(palmitoyloxy)-propyl]-(R)-cysteinyl-(lysyl) 3-lysine (Pam2 CSK4) and ODN M362 were reconstituted in endotoxin-free water, then diluted to the desired concentration in sterile PBS. For in vivo experiments, as previously described (Kirkpatrick et al, 2018; Ware et al, 2019), the indicated ligands were placed in an Aerotech II nebulizer driven by 10L/min air supplemented with 5% CO2 for 20 min. The nebulizer was connected by polyethylene tubing to a polyethylene exposure chamber. 24 h prior to infections, 10 ml of Pam2 (4 µM) and/or ODN (1 µM) was delivered via nebulization to unrestrained mice for 20 minutes, and then mice were returned to normal housing. For in vitro experiments, Pam2-ODN was added to the culture media 4 h prior to inoculation with virus.

### Reverse-Phase Protein Array

To simultaneously evaluate the expression of 161 regulatory proteins and phospho-proteins in HBEC-3kt cells after exposure to either PBS, Pam2, ODN or Pam2-ODN, a targeted high-throughput screening proteomic assay was performed by the Reverse Phase Protein Array Core Facility at The University of Texas MD Anderson Cancer Center (Tibes *et al*, 2006; Hennessy *et al*, 2010). The RPPA included 4 biological replicates per treatment condition, and data is available at GitHub (www.github.com/evanslaboratory/OBIF).

### Infection Models

For in vivo infections, frozen stock (2.8 × 107 50% tissue culture infective doses [TCID50] ml−1) of influenza A H3N2, virus was diluted 1:250 in 0.05% gelatin in Eagle’s minimal essential medium and delivered by aerosolization for 20 min to achieve a 90% lethal dose (LD90) to LD100 (∼100 TCID50 per mouse). Mouse health was followed for 21 d post infection. n = 15 mice per condition. Animals were weighed daily and sacrificed if they met euthanasia criteria, including signs of distress or loss of 20% pre-infection body weight. For in vitro infections, IAV (multiplicity of infection [MOI] of 1.0) was added to cells in submerged monolayer and viral burden was assessed 24 hours post infection.

### Pathogen burden quantification

To measure transcript levels of IAV nucleoprotein (NP) gene, samples were harvested in RNAlater and RNA was extracted using the RNeasy extraction kit. 500 ng total RNA was reverse transcribed to cDNA by using an iScript cDNA synthesis kit and submitted to quantitative reverse transcription-PCR (RT-PCR) analysis with SYBR green PCR master mix on an Bio-Rad CFX Connect Real-Time PCR Detection System. Host 18S rRNA was similarly probed to determine relative expression of viral transcripts.

### Omics Dataset Formatting

OBIF’s input in R requires an analysis-ready data matrix m with expression values and of dimensions f × n, where f is the number of features as rows and n is the number of samples S as columns. The appropriate sample order in dimensions n of m is:

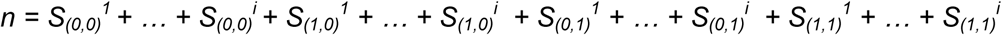

The subscripts denote the condition of the samples: exposed to neither factor (0,0), exposed to factor A alone (1,0), exposed to factor B alone (0,1) or exposed to both factors A and B (1,1). The superscripts represent the sample replicates from 1 to i within each of the four conditions.

To improve detection of interaction effects, OBIF allows sequential transformation of an unscaled original data matrix with background correction, log2-transformation, quantile normalization or a combination of these if needed. Background correction reduces noise to signal ratio at the lower limits of detection and methods vary per platform with code extensions are available at GitHub for microarray data using the lumi package, and for count-based sequencing data using rpm, rpkm, fpkm and tpm thresholds. Log2-transformation of continuous and count-base data is incorporated to provide a Gaussian-like data distribution, and quantile normalization is used to minimize the variance between samples during data scaling (Lo *et al*, 2015; Abrams *et al*, 2019) with OBIF to meet the statistical assumptions needed for two-way ANOVA analysis of interaction terms in a dataset (Slinker, 1998; Foucquier *et al*, 2015).

### Interaction analysis

To evaluate significant interaction terms between factors at the whole “-ome” level, OBIF performs a multiple linear regression across the expression values in a dataset:

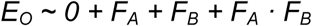

where the interaction analysis of the Omics expression levels (E_O_) is equivalent to a two-way ANOVA analysis where the intercept is referenced to the control samples (0) and returns a statistical summary of terms for the individual factor A (F_A_), factor B (F_B_) and their interaction (F_A_ · F_B_). Goodness of fit is calculated from the adjusted R^2^ values, and overall significance is determined by the p-values of F-statistics of the regression. Unscaled original data and scaled data with OBIF are compared to evaluate improvement in detection of significant interaction terms in a given dataset.

### Full Factorial Analysis

#### Expression Analysis

To perform differential expression analysis for detection of DEMs, OBIF fits a fixed-effects model to the expression data of each feature:

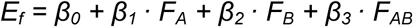

where the expression level of features (E_f_) is a function of the estimated β coefficients for the main effects of individual factor A (F_A_) and factor B (F_B_) and their combination (F_AB_). After regression, empirical Bayesian shrinkage of the standard errors is used to stabilize inferences of t-statistics, F-statistics, and log-odds used for differential expression analysis. Q-values are then calculated using the Benjamini and Hochberg method to reduce the false discovery rate (FDR). Alternatively, code extensions for are available at GitHub to perform Bonferroni corrections or calculate Tukey Honest Significant Differences adjustment for multiple testing instead of FDR.

#### Contrast analysis

To analyze the remaining factorial effects in the fitted linear model of feature expression, the coefficients and standard errors will be estimated typical of a two-way ANOVA from a set of contrasts that define the SMEs of each factor and their interaction effect:

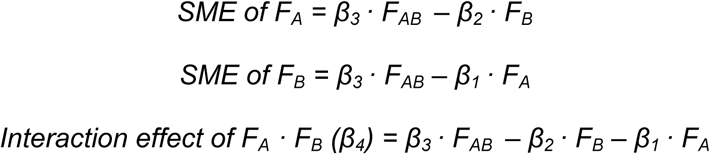

The standard errors calculated use a significance threshold (p-value < 0.05) to determine if DEMs with F_AB_ (Pam2-ODN) are susceptible to SME or interaction effects. Selection of iDEMs is based on DEMs of F_AB_ with a significant interaction effect.

### Mixed-effects model

To evaluate performance of full factorial analysis with OBIF, detection of interaction effects at the level of individual features is compared to a mixed-effect model (Caetano *et al*, 2018):

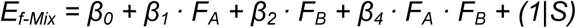

where the expression level of features in a mixed-effect model (E_f-Mix_) is a function of the estimated β coefficients for the fixed effects of individual factor A (F_A_) and factor B (F_B_) and their interaction (F_A_ · F_B_) with a random effect (1|S) for all sample conditions (*S*_*(0,0)*_, *S*_*(1,0)*_, *S*_*(0,1)*_, *S*_*(1,1)*_). After regression, empirical Bayesian shrinkage of the standard errors is used to stabilize inferences of t-statistics and F-statistics. The standard errors calculated from the interaction term use a significance threshold (p-value < 0.05) to determine significant interaction effects.

### Beta-uniform mixture model

Interaction p-values are extracted from the interaction term of mixed-effects model and from the interaction effect contrast of full factorial analysis. Independently, a beta-uniform mixture model is fitted to these sets of p-values (Pounds *et al*, 2003; Ji *et al*, 2005) to compare their discrimination ability using their receiver operating characteristic area under the curve (ROC AUC). Using the beta-uniform mixture models, we calculated the number of true positives (TP), false positive (FP) and false negatives (FN) detections (Pounds *et al*, 2003; Zhang *et al*, 2012) at the threshold level of iDEM selection (interaction p-value = 0.05) to estimate their precision and recall proportion:

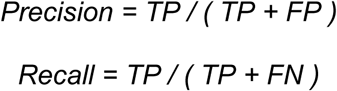

### Unsupervised clustering of features with OBIF

#### Hierarchical clustering and heatmaps of DEMs

All DEMs were represented in heatmaps after hierarchically clustering using Ward’s minimum variance method with Euclidean distances of log_2_FC values to compute dissimilarity by rows (features) and by columns (samples). Column dendrograms were plotted to represent the distance between samples, vertical side bar colors summarize DEMs according to their and horizontal side bars colors represent sample types by factors. Color scale keys indicate the levels of feature expression with upregulation in red and downregulation in green.

#### Principal component analysis of expression profiles

DEMs with F_AB_ (Pam2-ODN) were clustered by principal component analysis based on the mean linear fold change difference to reveal the expression patterns biologically present across all factors FA, FB and FAB. Principal components 1 and 2 were used for plotting DEMs with FAB and the variability between features is marked in each axis. EPs were identified in the clusters for each individual feature according to Table 1.

### Enrichment analysis

To provide biological interpretation of the full factorial analysis and classification of features, enrichment analysis was integrated in the pipeline to determine candidate effectors and regulators of synergy, biological pathways and functional processes. Sets for DEMs, EPs, DEMs with SMEs and iDEMs are uploaded independently, and enrichment analysis is performed with IPA software (QIAGEN, Hilden, Germany) for core analysis using the expression levels of features. Both gene and chemical Ingenuity Knowledge Base modules are used as enrichment reference, considering only experimentally observed confidence levels for identification of direct and indirect relationships. The thresholds of significance for canonical pathways, upstream analysis, diseases & functions, regulator effects and network analysis are ≥ 2 for activation z-score and < 5% false discovery rates for all predictions.

### DNA-binding ELISA

HBEC-3kt were grown to 80-100% confluence in 24-well plates and treated with PBS, Pam2, ODN, or Pam2-ODN for the indicated durations. Measurements of DNA-binding of members of NF-κB and AP-1 transcription factor members from whole cell lysates were made using their respective TransAM Kit according to product directions. For signal detection, samples were read immediately for absorbance at 450 nm with reference wavelength at 655 nm on a microplate reader. Experiments were repeated in triplicate and statistical analysis was performed with unpaired student’s t test using GraphPad Prism 8.0 with a significance threshold of p-value <0.05.

### Detection of Nuclear Translocation

#### Transcription factor staining and image acquisition

HBEC-3kt were grown to 80-100% confluence in 100mm dishes and treated for 30 min with PBS, Pam2, ODN or Pam2-ODN with or without pretreatment with NF-κB inhibitor IMD-0354 at 25 ng/μL for 16 h. Cells were detached from the plate with a 5 min incubation at 37 °C degrees with 3 ml of Accutase to prevent additional activation of transcriptional activity. Cells were pelleted in individual 15 ml tubes at 500 g for 5 min and suspended in 500 μL of eBioscience FOXP3 fixation/permeabilization buffer for 15 min at room temperature. Cells were stained with a LIVE/DEAD Fixable Near IR Dead Cell Dye and with a 1:1000 dilution of NF-κB p50 (E-10) Alexa Fluor 647, NF-κB p65 (F-6) Alexa Fluor 488, c-Jun (G-4) Alexa Fluor 594 and c-Fos (D-1) Alexa Fluor 546 conjugated antibodies for 1 h on ice and protected from light. After incubation, cells were pelleted and washed with 200 μL of sterile PBS 4 times, then resuspended in 100 μL sterile PBS. After the last wash, cells were pelleted and resuspended in 50 μL of sterile PBS and nuclear DAPI staining at 0.5 μg/mL was performed just prior to data acquisition on ImageStreamX MII.

#### Data acquisition with ImageStreamX MKII

HBEC-3kt images were acquired using INSPIRE software on the ImagestreamX Mark II imaging flow cytometer (Amnis Corporation) at 40× magnification, with lasers 405 nm (85.00 mW), 488 nm (200.00 mW), and side scatter (782 nm) (1.14 mW). 10,000 images per sample acquired include a brightfield image (Channel 1 and 9), p65 Alexa Fluor 488 (Channel 2), c-Fos Alexa Fluor 546 (Channel 3), c-Fos Alexa Fluor 594 (Channel 4), side scatter (Channel 6), DAPI (Channel 7), and p50 Alexa Fluor 647 (Channel 3). The laser outputs prevented saturation of pixels in the relevant detection channels as monitored by the corresponding Raw Max Pixel features during acquisition. For image compensation, single color controls were stained with all fluorochromes and 500 events were recorded with each laser for individual controls. Fluorescent images were taken in all channels with brightfield LEDs and scatter lasers turned off to accurately capture fluorescence. Individual single-color control file was then merged to generate a compensation matrix and all sample files were processed with this matrix applied.

#### Nuclear translocation analysis

After compensation for spectral overlap based on single color controls, analysis was performed and individual cell images were created using IDEAS® software version 6.1. Cell populations were hierarchically gated first by single cells, then cells in focus, then negative selected for live cells, and finally as double positive for both DAPI and the transcription factor subunit of interest (Figure EV3B). The spatial relationship between the transcription factors and nuclear images was measured using the ‘Similarity’ feature in the IDEAS software to quantitate the mean similarity score in the cell populations per sample. A similarity score >1 represents nuclear translocation, and the shift in distribution of nuclear translocation between two samples was calculated using the Fisher’s Discriminant ratio (Rd value) (Maguire *et al*, 2015).

### Statistical Analysis

Statistical analyses were performed using Prism 8 (GraphPad, San Diego, CA) and R. Kaplan-Meier curves were used for survival analyses and logrank (Mantel-Cox test) was used for paired group comparisons. Analysis of viral NP expression was performed using a two-way ANOVA with post hoc Tukey analysis for paired comparisons that was adjusted for multiple testing. Analysis of DNA-binding activity *in vitro* was performed using a student’s t test for comparisons between 2 groups, or using one-way ANOVA for comparison between multiple groups. Grouped data is shown as means +/- standard error of the mean, with experiments with *n* < 5 showing individual sample values. To verify the statistical assumptions for each test, Gaussian distribution was evaluated with Saphiro-Wilk test, and equal variance between two samples was evaluated with F-tests, or for more than two samples with Barlett’s or Levene’s test. Simultaneous multiple outlier detection was performed using the robust regression and outlier removal (ROUT) method with a q value of 5% (maximum FDR). Treatment allocation of animals was randomized in the experiments, though assessment could not be blinded. A pre-specified minimum requirement of 3 biological replicates for *in vitro* studies and 10 for *in vivo* studies.

### Data Availability

The data and code in this study are available in the following databases:

- OBIF R Package: GitHub Evanslaboratory/OBIF (www.github.com/evanslaboratory/OBIF)
- OBIF R Code: GitHub Evanslaboratory/Extensions (www.github.com/evanslaboratory/Extensions)
- Microarray data: Gene Expression Omnibus GSE28994 (https://www.ncbi.nlm.nih.gov/geo/query/acc.cgi?acc=GSE28994)
- RNA-seq data: Gene Expression Omnibus GSE109000 (https://www.ncbi.nlm.nih.gov/geo/query/acc.cgi?acc=GSE109000)
- Reverse-phase protein array data: GitHub Evanslaboratory/Datasource (www.github.com/evanslaboratory/Datasource)
- Reverse-phase protein array data: EMBO Molecular Medicine DOI:10.15252/emmm.201809960 (https://www.embopress.org/doi/abs/10.15252/emmm.201809960)
- Metabolomics data: Frontiers in Pharmacology DOI:10.3389/fphar.2019.00754 (https://www.frontiersin.org/articles/10.3389/fphar.2019.00754/full)

## Acknowledgements

The research reported here was supported by The University of Texas System and Mexico’s Consejo Nacional de Ciencia Y Tecnología (CONACYT) through the ConTex Postdoctoral Fellowship Program to J.P.G., by NIH grants R01 HL117976, DP2 HL123229 and R35 HL144805 to S.E.E. and by P30CA016672 to MD Anderson Cancer Center. The opinions expressed are those of the authors and do not represent views of these funding agencies. The Functional Proteomics RPPA Core facility is supported by MD Anderson Cancer Center Support Grant # 5 P30 CA016672-40. The Advanced Cytometry & Sorting Core Facility is supported by NCI P30CA016672 and is equipped for Imaging Flow Cytometry at MD Anderson Cancer Center. J.P.G. acknowledges and thanks the Methods in Epidemiologic, Clinical and Operations Research (MECOR) Program from the American Thoracic Society (ATS) and Asociación Latinoamericana de Tórax (ALAT) for their support and dedication in building research capacity in Latin America and other countries around the globe, particularly to their faculty Dr. Fernando Holguin and Dr. Altay Souza for their critical feedback.

## Author contributions

Conforming to the ICMJE criteria, all authors gave approval of the final version to be published and contributed to writing or revising the article critically for important intellectual content. Conforming to the CRediT criteria: J.P.G. and S.E.E. were involved in conceptualization of the project, visualization and writing of original draft, review & editing, and funding acquisition; J.P.G. was involved in data curation, formal analysis and methodology by conceiving, implementing and validating the systems model; J.P.G., V.V.K., T.C.R. and S.J.W. were involved in investigation by performing in vitro and in vivo experiments; J.P.G., V.V.K., T.C.R., S.W., S.J.W., J.Z., J.W., Y.W. and S.E.E. were involved in supervision by interpreting results; S.W., R.S., M.S.C., S.J.M., and F.M.J. were involved in providing resources by generating datasets for analysis and validation of the model; J.PG., J.Z. and J.W. were involved in formal analysis, software and validation by performing bioinformatics analyses; Y.W. and S.E.E. were involved in supervision, project administration and validation of the project.

## Expanded View Figure legends

**Figure EV1.**
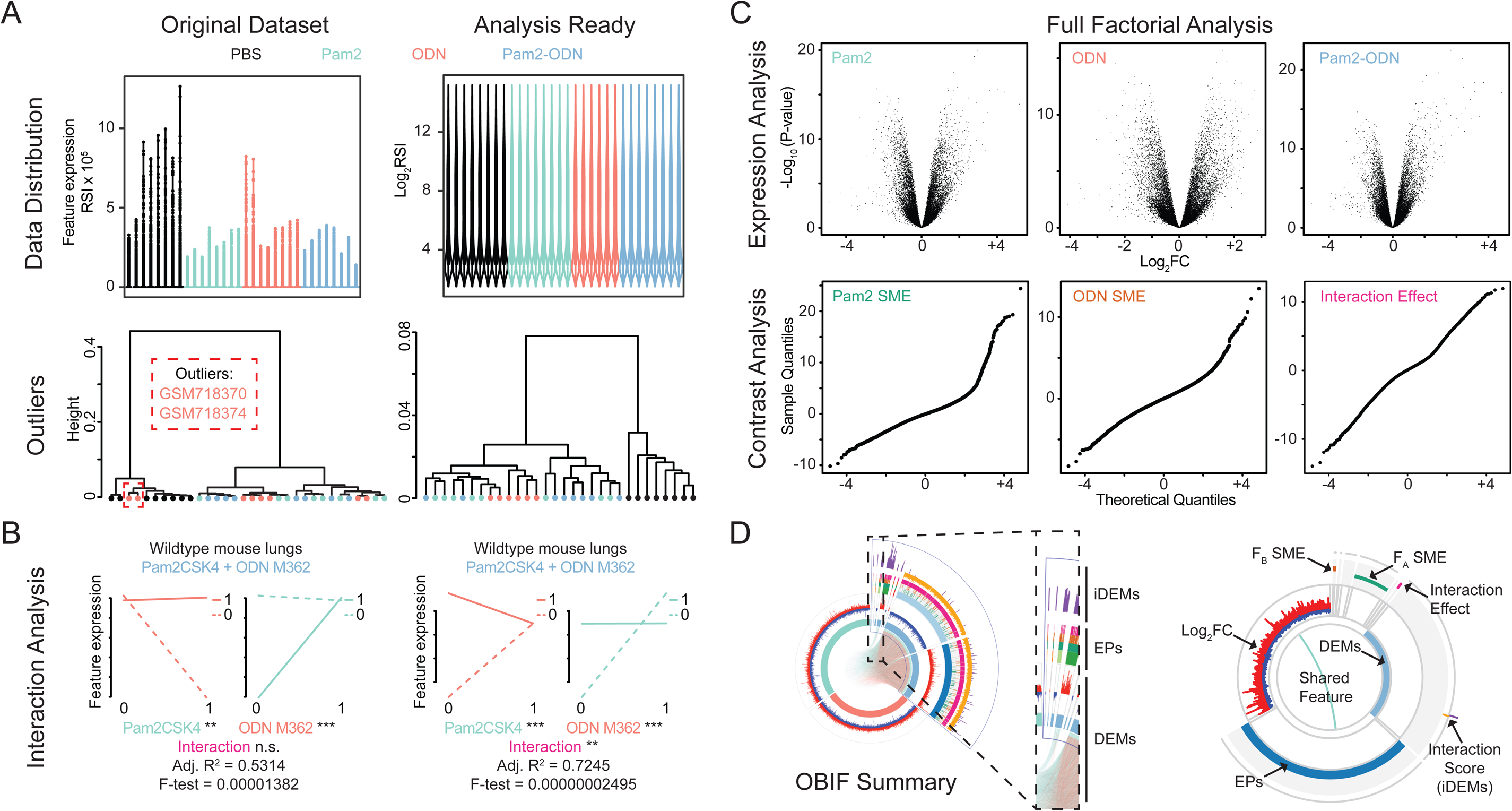
Overview of model fitting of Omics datasets during analysis with OBIF. (A) Quality control plots assess data distribution of GSE28994 with violin plots (*top*) and detect potential outliers by hierarchical clustering (*bottom*) in both the pre-processed original dataset (*left*) and the analysis-ready dataset (*right*). (B) Interaction analysis of factorial effects at the whole transcriptome level, demonstrating interaction plots, coefficient significance and goodness of fit per platform. (C) Representative volcano plots of full factorial analysis from analysis ready data for each condition after expression analysis (*top*) and Q-Q plots of moderated t-statistics for each multi-factor effect after contrast analysis (*bottom*). (D) Visual summary of OBIF’s outputs including DEMs, EPs and iDEMs (*left*) plotted into 3 rings (*right*): (i) DEMs, where inner links represent shared features between DEMs followed by their log_2_FC values; (ii) EPs, where inner sectors represent individual profiles (I to VIII) followed by their of significant F_A_-SME (green), F_B_-SME (orange) or F_A_·F_B_ interaction effect (pink); and (iii) iDEMs, represented by their synergistic or antagonistic interaction scores. FC, fold change; DEMs, differentially expressed molecules; EPs, expression profiles; iDEMs, interacting DEMs; SME, simple main effect.

**Figure EV2.**
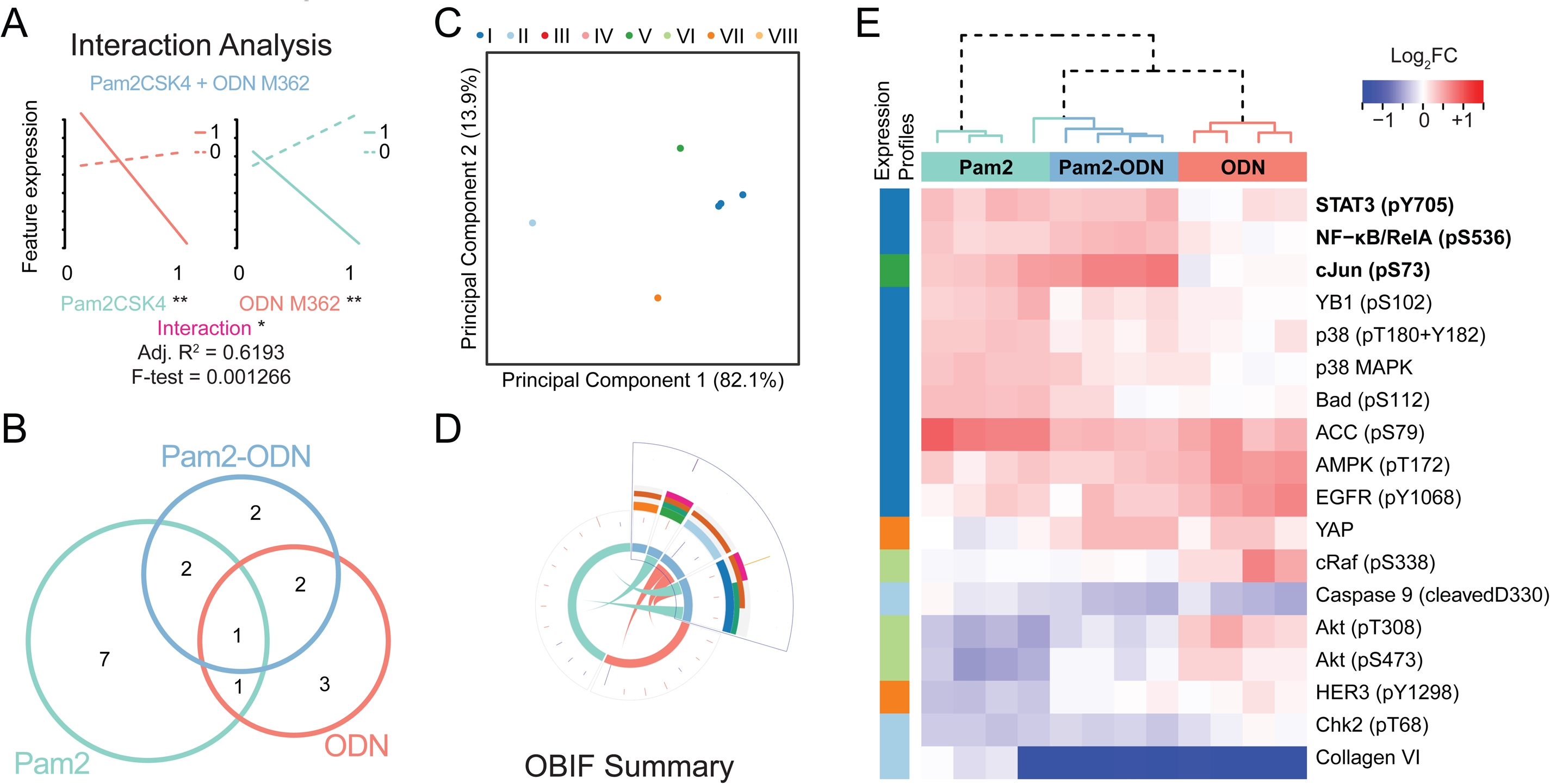
OBIF analysis of RPPA data from HBEC-3kt treated with single or dual ligands. (A) Interaction analysis of factorial effects at the whole proteome level, demonstrating interaction plots, coefficient significance and quality of model fitness per platform. (B) Euler diagram of DEMs identified in A. (C) Principal component analysis of dual factor DEMs in B clustered by EPs. (D) OBIF summary of molecular drivers of synergy in B-C. (E) Heatmap of expression values of DEMs in B with expression profiles shown per feature (rows). FC, fold change; DEMs, differentially expressed molecules; EPs, expression profiles.

**Figure EV3.**
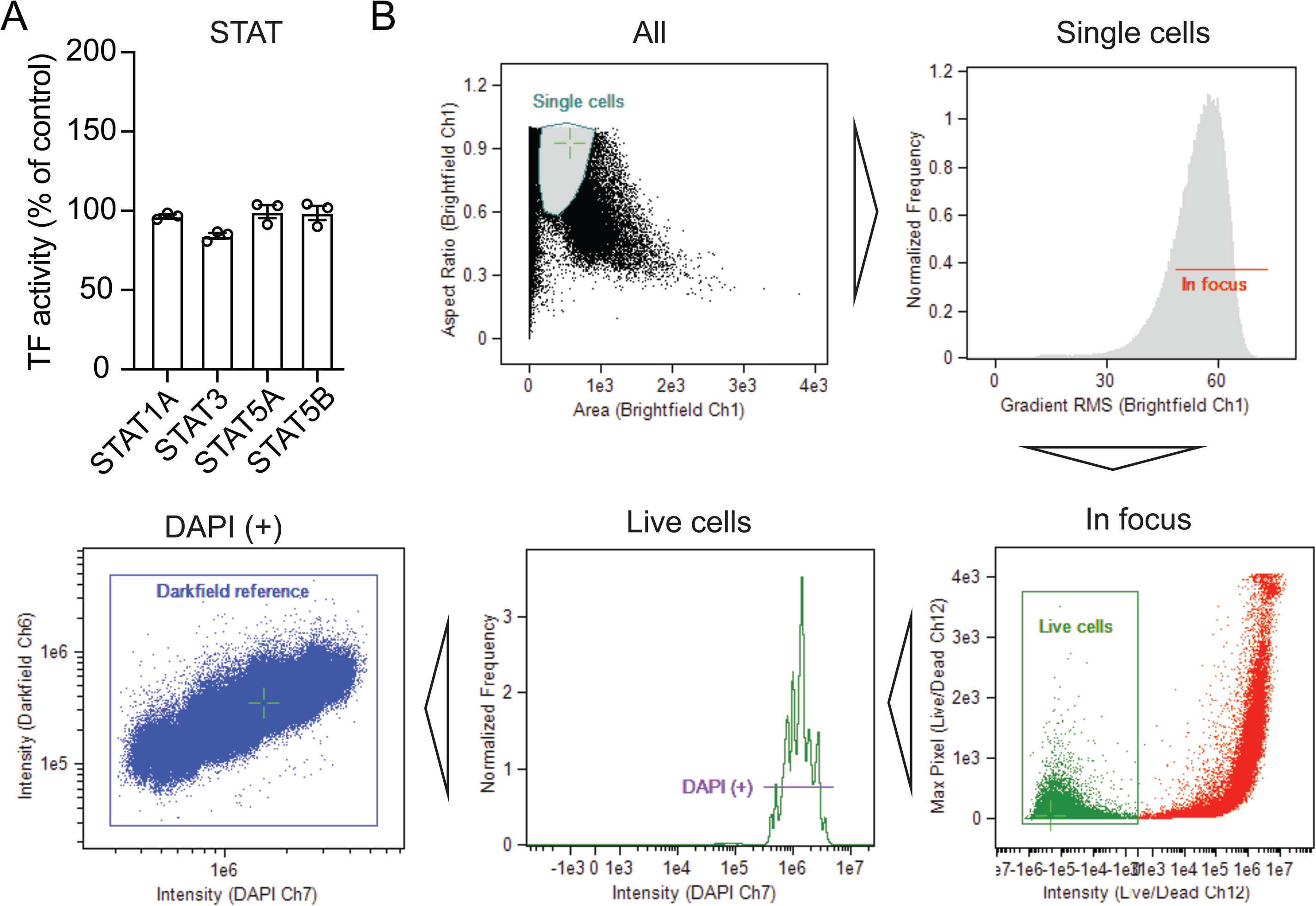
STAT family data and gating strategy for imaging flow cytometry. (A) Transcription factor activity of STAT subunits 15 min after treatment of human lung epithelial cells with Pam2-ODN. n = 3 samples/condition. (B) Gating strategy used during single cell imaging flow cytometry for simultaneous assessment of all transcriptional subunits.

**Figure EV4.**
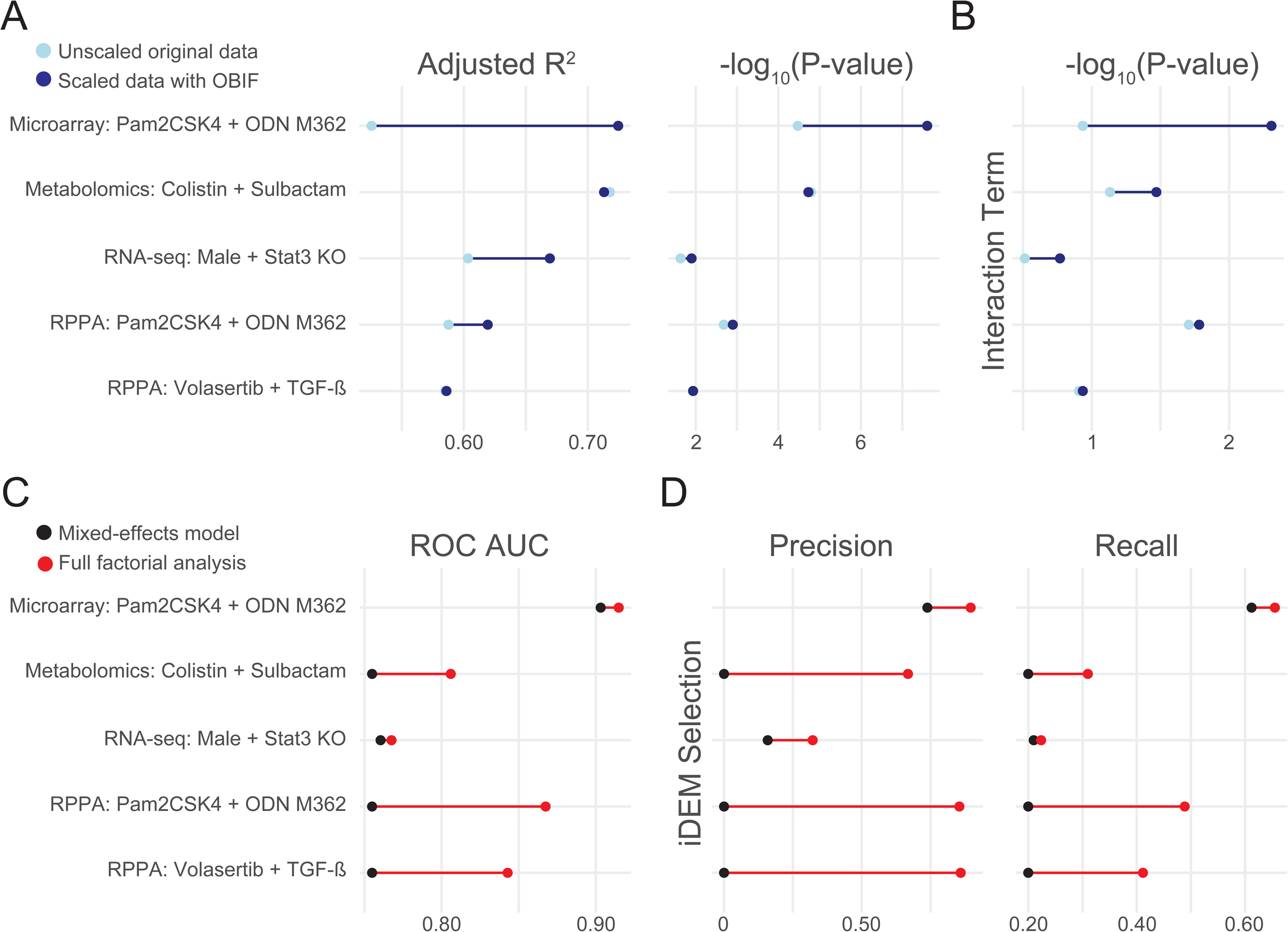
OBIF improves detection of interaction effects across platforms and factor classes. (A) Comparative performance of data scaling during interaction analysis at the whole “-ome” level showing overall fitness and significance of two-way ANOVA. (B) Significance level of interaction term between factors detected in A. (C) Comparative performance of statistical methods to detect interaction effects at the individual feature level using a beta-uniform mixture model of interaction p-values. (D) Precision and recall fractions at iDEM selection threshold calculated from C. OBIF, Omics-based interaction framework; ROC, receiver operating characteristic; AUC, area under the curve.

